# GPU-accelerated homology search with MMseqs2

**DOI:** 10.1101/2024.11.13.623350

**Authors:** Felix Kallenborn, Alejandro Chacon, Christian Hundt, Hassan Sirelkhatim, Kieran Didi, Sooyoung Cha, Christian Dallago, Milot Mirdita, Bertil Schmidt, Martin Steinegger

**Affiliations:** Department of Computer Science, Johannes Gutenberg University Mainz, Mainz, Germany; NVIDIA Corp; School of Biological Sciences, Seoul National University, Seoul, Republic of Korea; Interdisciplinary Program in Bioinformatics, Seoul National University, Seoul, Republic of Korea; Department of Biostatistics and Bioinformatics, Duke University, Durham, NC 27705, United States; Department of Cell Biology, Duke University, Durham, NC 27705, United States; Institute of Molecular Biology and Genetics, Seoul National University, Seoul, Republic of Korea; Artificial Intelligence Institute, Seoul National University, Seoul, Republic of Korea

## Abstract

Rapidly growing protein databases demand faster sensitive sequence similarity detection. We present GPU-accelerated search utilizing intra-query parallelization delivering 6x faster single-protein searches compared to state-of-the-art CPU methods on 2×64 cores—speeds previously requiring large protein batches. It is most cost effective, including in large-batches at 0.45x MMseqs2-CPU speed (8 GPUs delivering 2.4x). It accelerates ColabFold structure prediction 31.8x compared to AlphaFold2 and Foldseek search 4-27x. MMseqs2-GPU is open-source at mmseqs.com.

---

Many advances in biology have been enabled by computational tools retrieving evolutionary related sequences (homologs) from reference databases (1–4). Building on the sequence-based protein homology paradigm (5, 6), these tools detect homologs to an input query among millions to billions of reference entries by searching for similar amino acid sequences. Homology search is critical for inference of protein properties (7–9) such as early secondary structure prediction (10). Remote homologs have been shown to be pivotal as input to contemporary deep learning methods like AlphaFold2 and others (11–13) to predict accurate 3D structures (14–16). To retrieve remote homologs, sensitive tools to detect pairwise similarity between a query and reference sequences in a database are required. Theoretically, high sensitivity can be achieved by applying the dynamic programming-based, gapped Smith-Waterman-Gotoh algorithm (17, 18) to find the optimal path (alignment) (19) for each query-reference alignment-matrix. However, the ever-growing size of reference sequence databases (15) renders this exhaustive approach impractically slow. As a result, heuristic-based methods like BLAST (1), PSI-BLAST (20), MMseqs2 (ref. 4), and DIAMOND (3) incorporate filtering techniques to prune the majority of dissimilar sequences before executing the computationally expensive gapped computation. This is typically done by employing a seed-and-extend strategy, in which short k-mer words (“seeds”) are indexed and matched, followed by their extension to gapped alignments. Sensitive aligners like HMMER (2) and HHblits (21) instead apply a simplified dynamic programming filter, which scores all gap-free paths (strict diagonals) of the alignment-matrix between sequence pairs to find the highest scoring gapless match. Unlike k-mer-based methods, being a lower-bound approximation of a gapped alignment, gapless alignments result in a score for all pairs at the cost of computational efficiency. Several approaches were explored to achieve higher execution speed regardless of heuristic, like central processing unit (CPU) specific instructions and parallelization, or employing specialized hardware like Field-Programmable Gate Arrays (FPGAs; 22), and Graphics Processing Units (GPUs; 23, 24).

Here, we present two novel GPU-accelerated algorithms (Fig. 1a,b) for gapless filtering and gapped alignment using position-specific scoring matrices (PSSMs; 25). Integrated in MMseqs2, they achieve speed comparable to fast k-mer-based approaches while maintaining high sensitivity. We confirm these findings through two benchmarks, one focusing on homology retrieval, in the best setting achieving 17-fold (8 GPUs) or six-fold (1 GPU) speed-up compared to next fastest CPU-based method BLAST, and on query-centered multiple sequence alignment (MSA) generation for structure prediction, achieving 176-fold speed-up compared to JackHMMER and HHblits in AlphaFold2.

**Fig. 1.**
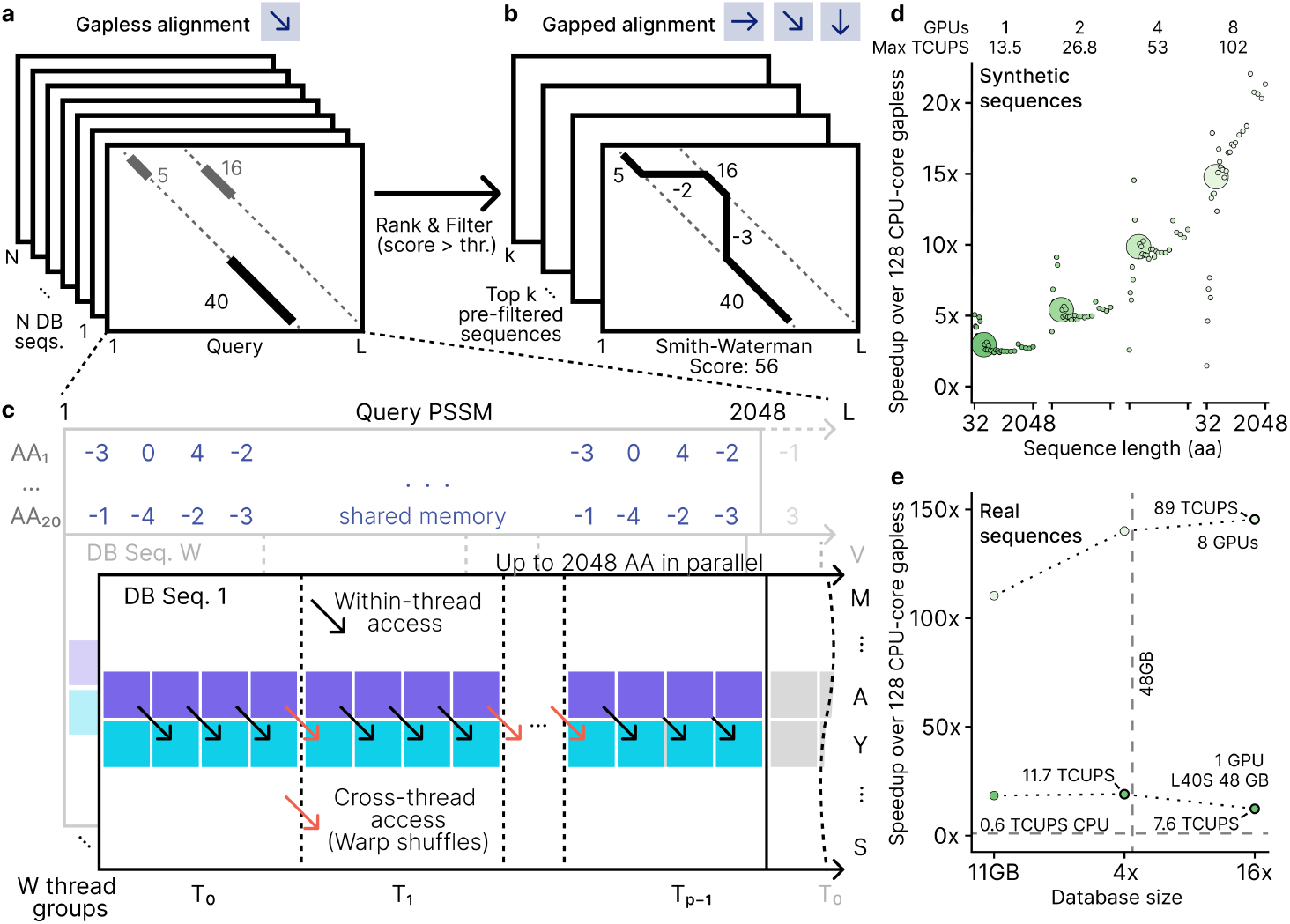
MMseqs2-GPU workflow and gapless alignment performance. (a) The gapless alignment involves scanning database sequences against the query, followed by ranking and filtering based on alignment scores. Only sequences with scores above a threshold proceed to (b) gapped alignment using the Smith-Waterman-Gotoh algorithm. In the GPU-optimized gapless alignment (c), the query profile is split into segments of up to 2048 amino acids, which are loaded into shared memory for fast access across thread groups, avoiding slower global memory access. Each thread computes alignment scores independently within its segment, while cross-thread warp shuffles enable efficient data sharing for diagonal dependencies. (d) Speedup of 1, 2, 4 and 8 GPU execution of the gapless kernel on randomly generated sequence pairs of lengths 32 to 2048 in comparison to 128-core CPU execution. (e) Speedup of 1 and 8 GPU executions relative to a 128-core CPU system for 6370 queries against target sets of 1x, 4x, and 16x a 30M protein database (Methods “Sensitivity”). 16x exceeds single GPU RAM and is processed with database streaming with 7.575TCUPS/11.676TCUPS ≈ 64.9% of in-memory processing speed.

Our GPU-accelerated gapless filter maps query PSSMs to columns and reference sequences to rows in a matrix, then processes each matrix row in parallel, while utilizing shared GPU memory to optimize access to PSSMs (Fig. 1c) and 16-bit floating-point numbers packed in a 32-bit word (half2) to maximize throughput. For gapped alignment, we incorporated a modified CUDASW++4.0 (ref. 26) that operates on PSSMs, employing a wavefront pattern to efficiently handle dynamic programming dependencies.

In a synthetic benchmark of random amino acid sequences, the gapless GPU kernel achieved up to 2.8x (single L40S) and 21.4x (eight L40S) speed-ups and peak performance of 13.5 TCUPS and 102 TCUPS, respectively, compared to a 2×64 CPU-core server (Fig. 1d), outperforming previous acceleration methods by one-to-two orders of magnitude (22, 23). When measured on real protein sequences, relative speedup compared to CPU increased to 18.4x and 110x, while TCUPS peaked at 11.3 and 67.5 for 1 and 8 GPUs, respectively (Fig. 1e; Methods, “Database scaling benchmark”; Supplementary Data 1, “Ungapped peak performance”). It outperforms previous accelerator-based approaches for gapless filtering by one-to-two orders of magnitude, exceeding their maximum reported performances of 1.7 TCUPS on an Alveo U50 FPGA (22) and 0.4 TCUPS on a K40 GPU (23).

Prior to evaluating homology search speed of the combined gapless and gapped alignment workflow, we sought parameters allowing to reach similar sensitivity as measured on the MMseqs2 benchmark (4). This resulted in evaluated methods reaching ROC1 between 0.391-0.409, specifically setting MMseqs2-CPU k-mer with the `-s 8.5` setting and DIAMOND with the `--ultra-sensitive` option. Furthermore, to examine the ability of MMseqs2-GPU to increase sensitivity, we investigated iterative profile searches, where initial hits are converted into PSSMs that encode probabilities for each residue in a sequence. MMseqs2-GPU reached ROC1 of 0.612 and 0.669 at two and three iterations, respectively. This was slightly higher than PSI-BLAST, reaching 0.574 and 0.591, and slightly lower than JackHMMER, which benefited from its Hidden-Markov-Model-based profiles in higher search iterations and reached 0.614 and 0.685 at two and three iterations, respectively (Methods, “Sensitivity”; Supplementary Data 1, “Sensitivity benchmark”).

We then benchmarked speed for homology search focusing on two common scenarios: a single query protein against a target database of roughly 30M sequences (single-batch), common for scientists working on a protein system, and a set of query proteins against the same 30M target database (large-batch, i.e. 6370 queries), common for proteome analysis. MMseqs2-GPU on a single NVIDIA L40S outperformed the commonly-used JackHMMER, being 177 (single-) and 199 (large-batch) times faster (Fig. 2a; Supplementary Data 1, “Search tool comparison”). In single-batch, MMseqs2-GPU was 6.4 times faster than the second-fastest method BLAST (Fig. 2a, left), while at batch size 100 it was 1.9 as fast as the second-fastest in that setting, MMseqs2-CPU k-mer (Fig. 2a, middle). At batch size 6370, MMseqs2-CPU k-mer on a 128-core CPU system was about 2.2x faster than MMseqs2-GPU on a single L40S, however on an 8 GPU system, MMseqs2-GPU takes the lead at 2.4x the speed of MMseqs2-CPU k-mer (Fig. 2a, right). On a single H100 GPU system MMseqs2-GPU followed similar trends to one NVIDIA L40S while an A100 is 1.6x slower (Supplementary Data 1).

**Fig. 2.**
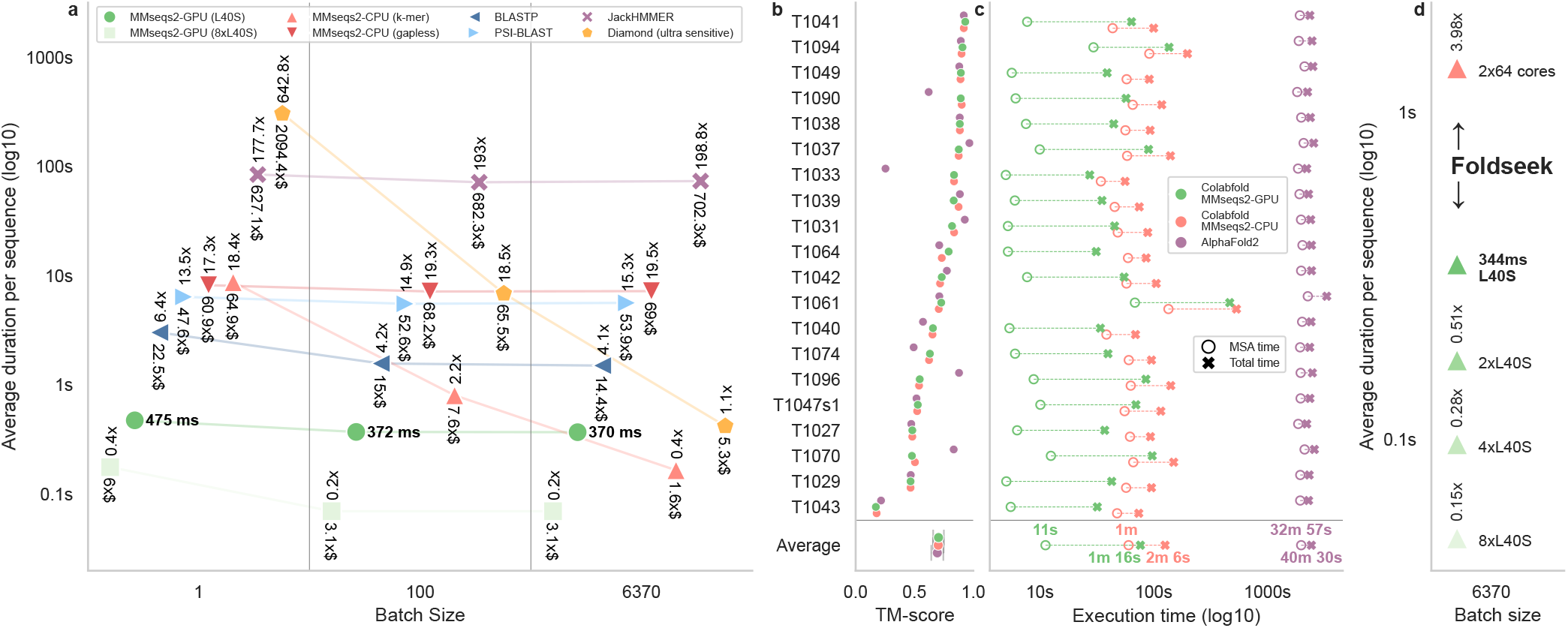
MMseqs2-GPU runtimes for homology search. (a) In single-batch processing, MMseqs2-GPU on a single L40S GPU (dark green; baseline execution time in bold, horizontal) is ∼six times faster than the second fastest BLAST (dark blue; top vertical numbers), and ∼178 times faster than JackHMMER (purple) on average for 6370 queries against a 30M target database. MMseqs2-GPU can split the target database across GPUs to enable larger database searches, also achieving a speedup compared to single GPU execution (up to five times faster on eight L40S compared to one; bright green vs. dark green). When present, JackHMMER values refer to measuring 10% of the queries, as execution time for 6370 queries was prohibitively inefficient. Similarly, execution time for Diamond ultra sensitive in the single-batch setting refers to 10% of the query set. For all batch sizes, MMseqs2-GPU on one L40S provides the lowest AWS cost for execution. In fact, even while MMseqs2-CPU k-mer in batch 6370 provides faster execution, it results 1.6x more costly than MMseqs2-GPU on one L40S (bottom vertical numbers). (b,c) Faster folding speeds at no accuracy cost. On 20 CASP14 targets, ColabFold leveraging MMseqs2-GPU (green) results 1.65 and 31.78 times faster than ColabFold using MMseqs2-CPU k-mer (orange) or AlphaFold2 using JackHMMER and HHblits (violet), respectively. All methods reach similar TM score (∼0.70±0.05). Here, MMseqs2 searched through 238M cluster representatives and subsequently expands to 1B members, while JackHMMER searched through 426M sequences and HHblits searched through 81M profiles containing 2.1B members. (d) Foldseek built on MMseqs2-GPU on one L40S (dark green; baseline execution time bold, horizontal) reaches four times the speed of Foldseek-CPU k-mer (orange; vertical numbers) even at large-batch size (6370) due to required higher sensitivity. Scaling to eight L40S GPUs speeds-up execution seven and 27 times compared to one L40S and Foldseek-CPU k-mer, respectively.

Diamond benefited from batching more than MMseqs2-GPU (Fig. 2a, slopes from left to right). Therefore, we further investigated Diamond’s batching behavior by using larger batch sizes (10,000 and 100,000 sequences). We found that Diamond’s average per-query speed does not improve and remains around 0.42s, which was measured on batch 6370. At batch size 6370, MMseqs2-GPU and MMseqs2-CPU k-mer reached 0.37 and 0.17 seconds per query, respectively, making them overall faster than Diamond (Methods, “Diamond scaling”; Supplementary Data 1, “Search tool comparison”).

To provide cloud cost estimates, we compared AWS EC2 pricing. MMseqs2-GPU on a single L40S instance was the least-expensive option across all batch sizes. MMseqs2-CPU k-mer on 2×64 cores was 60.9 and 1.6 times more costly for single-batch and large-batch workloads, respectively (Fig. 2a). Additionally, we also investigated energy consumption in the single-batch scenario. MMseqs2-GPU achieved the highest energy efficiency in a hardware configuration with four L40S GPUs, where it achieved 80.7 and 2.1 times higher energy efficiency than JackHMMER and MMseqs2-CPU k-mer, respectively (Supplementary Data 1, “Energy consumption”). Yet higher energy efficiency of 95 (vs. JackHMMER) and 2.5 (vs. MMseqs2-CPU) times could be observed when running MMseqs2-GPU on a system with 16 CPU-cores and one L4 GPU (Supplementary Data 1, “Energy consumption”).

Over time, MMseqs2’s k-mer-based filtering was optimized to handle smaller input batch sizes, particularly for web servers (27) and to generate MSA input for ColabFold (28), however requiring substantial RAM (up to 2TB) for fast MSA generation within ColabFold. MMseqs2-GPU addresses this issue, requiring only one byte per residue in the target database compared to ∼ 7 bytes per residue required for MMseqs2-CPU k-mer, which can further be reduced by clustered searches against representative sequences followed by member realignment. On top, MMseqs2-GPU allows searching large reference databases by distributing them across multiple GPUs, resulting in additional speed-up, or by streaming them from host RAM at a slow-down. The latter enables processing databases larger than the available GPU memory, albeit at 63 to 65% of in-GPU-memory processing speed (Fig. 1e, Supp. Fig. 1, Methods “Database Streaming”).

Computational protein structure prediction has become ubiquitous since the release of AlphaFold2 (ref. 11) and RoseTTAFold (12) in 2021. Variations of these methods using alternative representations were introduced for higher throughput (29); however, methods that leverage query-centered MSAs remain the most accurate (14). We compared the homology search and structure prediction steps for the single batch scenario common for protein prediction servers for three variants: the canonical setting of AlphaFold2 leveraging HH-blits (21) and JackHMMER (2), and ColabFold leveraging either MMseqs2-CPU k-mer or MMseqs2-GPU on one L40S. The AlphaFold2 search pipeline targets 506M entries (426M sequences and 81M profiles, the latter containing ∼ 2.1B cluster members). The ColabFold-MMseqs2 pipeline conducts two three-iteration profile searches against in total 238M cluster representatives, and then expands and realigns to search one billion cluster members (Methods “Structure prediction”).

We compare these pipelines for structure prediction of 20 free modeling sequences from CASP14 (ref. 30). ColabFold utilizing MMseqs2-GPU was fastest, overall being 1.65x faster than ColabFold MMseqs2-CPU k-mer and 31.8x faster than AlphaFold2 leveraging HHblits and JackHMMER (Fig. 2c), showcasing MMseqs2-GPU’s suitability to metagenomics-scale searches. A key reason for this overall speedup is the MSA generation step: computing MSAs was 176.3x and 5.4x faster using MMseqs2-GPU compared to using JackHMMER+HHblits and MMseqs2-CPU k-mer, respectively. Consequently, while MSA computation on the CPU consumes the majority of runtime in AlphaFold2 (83% of total execution time; Supplementary Data 1, “Folding tool comparison”), often resulting in idle GPU time or necessitating splitting MSA and model inference onto different systems for optimal performance, ColabFold paired with MMseqs2-GPU crucially reduces MSA generation to only 14.7% of total execution time, allowing seamless execution on a single GPU-equipped system. These speed differences had no effect on prediction accuracy, with all three configurations achieving TM-score around 0.70±0.05 (Fig. 2b; Supplementary Data 1, “Folding tool comparison”).

MMseqs2 serves as the backbone to various methods through its modular architecture, such as the protein structure aligner Foldseek (31). Foldseek requires high sensitivity to obtain good structure alignment, reducing the advantage of fast k-mer-based double-diagonal search. We measured Foldseek’s search speed for CPU (Foldseek-CPU k-mer) and GPU (Foldseek-GPU) on 6370 protein structure queries sampled from the AlphaFold Database clustered at 50% structure identity, searching them against the same reference database. Foldseek-GPU on a single L40S outperformed Foldseek k-mer using 2×64 CPU-cores by a factor of 4, and was ∼27.3 times faster on eight L40S GPUs (Fig. 2d). Repeating the Foldseek SCOPe-based sensitivity benchmark with Foldseek-GPU yielded modest increases in sensitivity across Family (0.874 vs. 0.861), Superfamily (0.493 vs. 0.487), and Fold recognition (0.108 vs. 0.106) compared to Foldseek k-mer.

MMseqs2-GPU ranks as the fastest and cheapest evolutionary search tool across various experiments. Unlike word-based search methods (1, 3, 4), MMseqs2-GPU’s sensitive gapless filter, similar to JackHMMER, does not allow trading sensitivity for speed. However, reaching high-sensitivity through iterative profile searches is essential for detecting remote homologs, which is critical for tasks like structure prediction (14), making it an excellent choice for these workflows.

Our experiments ran on a server with 1TB RAM and 256 threads — high-performance computing resources typically out of reach for most academics. MMseqs2-GPU overcomes this limitation both by reducing memory requirements, and by offloading compute-intensive tasks to the GPU. Even on a cost-effective GPU like the NVIDIA L4 with 24GB RAM, which is available through platforms such as Google Colab Pro in combination with a 6-core/12-thread CPU and 64GB RAM, MMseqs2-GPU accelerates searches over CPU-based methods, offering a ten-fold speed increase over JackHMMER in searching UniRef90 (2022_01, 144M proteins, Methods & Supplementary Data 1 “Colab benchmark”). While GPU memory can be a limitation, MMseqs2-GPU mitigates this with four strategies: reduced memory footprint, efficient database streaming, partitioning, and clustered searches.

We demonstrated that MMseqs2-GPU reduces the end-to-end execution time of protein homology search and structure prediction without compromising accuracy, enabling higher throughput at lower cost (Fig. 2). Many bioinformatics applications rely on homology searches and could benefit immediately. For example, orthology inference tools that already use MMseqs2 can seamlessly switch to the GPU backend (32, 33), as well as structural protein search methods like Foldseek (31). Especially workflows that already leverage GPUs, but are impractically slow at inference, like retrieval augmented protein language models (34), stand to benefit. Therefore, MMseqs2-GPU broadens access to rapid and affordable homology searches.

## Online Methods

### A. Background: fast sequence comparison principles

Filtering is an essential step in homology search to reduce the overall execution time by ranking all reference database sequences quickly, and filtering a subset against which the computationally more demanding Smith-Waterman-Gotoh can be run. As one of the earliest tools, BLAST (1) proposed index structures for initial seed finding with bi-directional gapless extension to match a given minimum score.

MMseqs2 (4) introduced a filtering approach based on double-consecutive k-mer matches on the same diagonal. Through this approach, all k-mers of a reference database are stored in a random-access-memory (RAM) based index structure for quick retrieval (hereafter the index). For single query searches at high sensitivity, MMseqs2 generates long lists of similar k-mers from the query sequence and matches them against the index to check the filtering criterion of two consecutive matches on one diagonal. While highly optimized, random accesses to the index results in poor cache-locality.

Diamond (3) generates lists of k-mers for both queries and references in order to sort and compare them co-linearly. Instead of similar k-mers, Diamond utilizes multiple spaced-k-mer patterns. Co-linear comparison results in improved cache-locality at the expense of indexing and sorting overhead.

In contrast to word-based filtering approaches, HMMER (2) and HHblits (21) implemented a more sensitive, albeit slower ranking technique by simplifying the Smith-Waterman-Gotoh algorithm to perform a gapless alignment (i.e., excluding gaps). In essence, this allows finding the longest common subsequence (LCS) consisting of residue substitutions only between any pair of sequences, allowing for better resolution than word-based methods.

Modern hardware accelerators like GPUs lend themselves to highly parallel workflows through their high core count, albeit at lower operational complexity per core. Thus, the gapless filtering is well suited to exploit their capabilities, due to reduced instruction count with little data-dependencies, while additionally avoiding branching and random memory accesses typically employed in k-mer index lookups.

### B. MMseqs2-GPU algorithm and parallelization

Pairwise gapless alignments are computed between a query represented by the PSSM Q and each reference sequence. Q is constructed either through a single query sequence and a substitution matrix, or from a sequence profile, based on previous search results. Once computed, Q provides a score for placing any amino acid at any position along the length of the query. Next, our gapless filter computes a pairwise local alignment between the reference sequence S = (s1, …, s_n_) and the PSSM Q of length m by dynamic programming (DP), and populates a matrix M using the recurrence relation M [i, j] = max {M (i − 1, j − 1) + Q[i, s_j_], 0} for all 1 ≤ i ≤ m, 1 ≤ j ≤ n. Initialization is given by M [i, 0] = M [0, j] = 0. Algorithm 1 outputs the maximum value in M, which represents the score of the optimal local alignment without gaps (a.k.a., gapless) between S and Q. The top k sequences passing an inclusion threshold are then passed to a Smith-Waterman-Gotoh algorithm operating on profiles. Both of these algorithmic steps are executed on GPU.

#### Algorithm 1

Gapless filter score computation

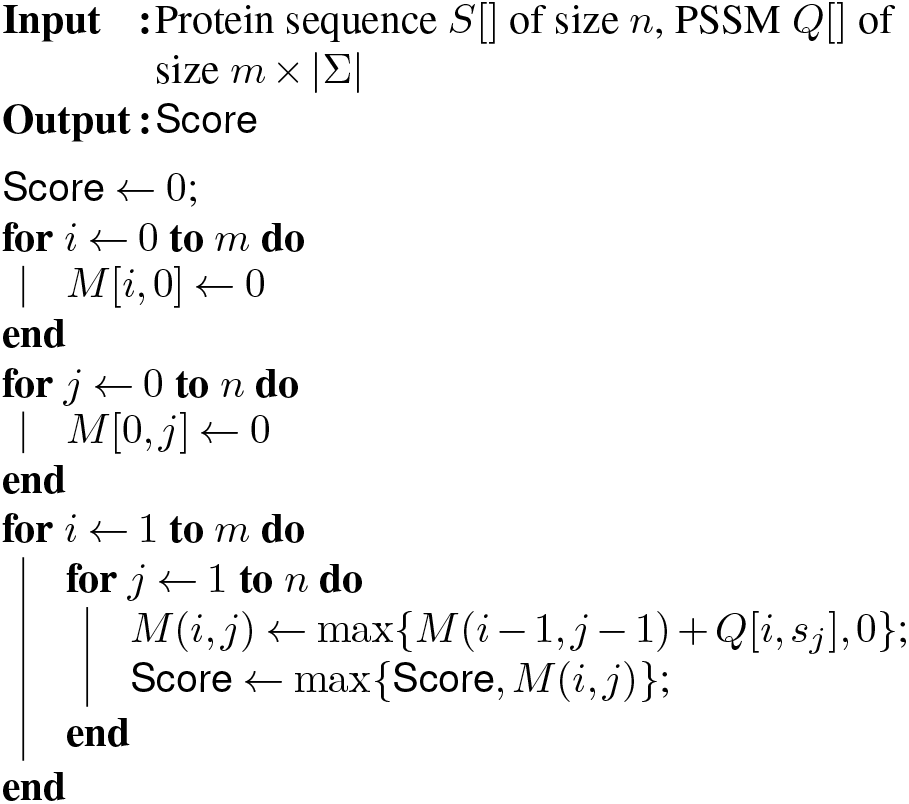

#### B.1. CPU-SIMD gapless filter

MMseqs2 offers a gapless filter accelerated on CPUs through Single Instruction Multiple Data (SIMD) instructions, through the `ungappedprefilter` module first introduced in MMseqs2 6.f5a1c. In this work, we describe the module, its integration into the MMseqs2 `search` workflow with `--prefilter-mode 1` and its extension to incorporate soft-masking of low-complexity regions with tantan (35)—in addition to the GPU integration described below. The filtering process begins by preparing a striped query profile (36) on a single thread, and finally utilizes all available CPU-cores to linearly compare all reference sequences on disk. During filtering, all soft-masked target database amino acids are represented as the unknown `X` residue.

At its core, the gapless alignment follows Farrar’s (36) striped Smith-Waterman-Gotoh first used in HMMER (2), which is adjusted to compute only the gapless diagonal. It avoids affine gap computation and requires only five SIMD operations to update a striped query segment to retrieve the maximum gapless diagonal score.

The algorithm uses 8-bit integers to represent alignment scores to maximize the parallel SIMD register usage. Scores that exceed the 8-bit limit of 255 are clamped to 255, which is significant and indicates a strong potential match. Such matches are then aligned using full Smith-Waterman-Gotoh. Depending on the supported instruction set of the CPU this gapless implementation uses 128-bit (SSE2) or 256-bit (AVX2) vector size. We also evaluated AVX512 and found only a marginal performance benefit and did not implement it. In addition, our implementation exploits multi-core CPUs by computing different target sequences independently using multi-threading.

#### B.2. GPU-accelerated gapless filter

##### Core algorithm

We present a novel implementation of a gapless filter on GPUs, designed to leverage the simplified dependency scheme inherent in gapless alignments for high parallelism and performance. Unlike the Smith-Waterman-Gotoh algorithm, where each cell in the DP matrix depends on its top, left, and diagonal neighbors, gapless alignment reduces this dependency to only the diagonal neighbor (Fig. 1). This crucial simplification enables parallel computation across all cells within the same row, potentially enhancing processing speed in comparison to more complicated parallelization schemes, such as wavefront parallelization typically employed to accelerate gapped Smith-Waterman computation. However, to realize this potential, the design of an appropriate parallelization scheme is required that optimizes necessary memory accesses and inter-thread communication, while efficiently handling highly variable sequence lengths.

##### Memory accesses and data types

To optimize memory accesses and performance, the query PSSM is stored in shared memory, facilitating fast, multi-threaded access (Fig. 1c). Meanwhile, reference sequence residues, which map to the y-axis of the DP matrix, are stored in global memory, requiring one byte per residue. Our calculations employ 16-bit floating-point numbers or 16-bit integers, supported by the DPX instructions available with NVIDIA’s Hopper GPU architecture, packed into a 32-bit word using half2 or s16×2 data types.

##### Thread grouping and tile optimization

Drawing inspiration from CUDASW++4.0 (26), our implementation assigns each gapless alignment task to a thread group, with typical sizes of 4, 8, or 16 threads. This configuration allows multiple thread groups to handle different alignments concurrently. The DP matrix is processed row-by-row, with each row partitioned among threads that are responsible for up to 128 cells each. To optimize performance across various query lengths, we employ different tile sizes, which are determined by the product of the thread group size and the number of columns processed per thread. They are realized as template parameters, with the group size constrained to be a divisor of 32, and half the number of columns being a multiple of 4. Optimal tile size configurations for different query lengths were identified via a separate grid search program, minimizing out-of-bounds computations when the query length is less than the tile size. As the processing begins, reference sequence residues are loaded from global memory, and each thread accesses its corresponding values from the scoring profile. To boost throughput, we designed the memory access pattern so that each thread loads eight consecutive 16-bit scores from shared memory in a single instruction.

##### Shared memory bank conflict mitigation

We further optimized shared memory access for thread groups of size four to mitigate bank conflicts. Shared memory is organized into 32 four-byte banks, and we utilize two-byte values in the PSSM, meaning two columns are packed per four-byte word. The PSSM is arranged in shared memory such that the i-th four-byte column maps to memory bank i mod 32.

A warp-wide load from shared memory is broken up into one or more hardware transactions of size 128 bytes served by the 32 memory banks. Bank conflicts occur when multiple accesses within the same transaction target the same bank but different addresses, leading to serialization. This can arise when multiple thread groups within a warp access the same PSSM columns but different rows (corresponding to different residues in the reference sequence). For group sizes of eight or larger, this is not an issue, as each group loads 8 × 16 = 128 bytes, thus fitting a single transaction. However, with a group size of four, each group loads only 4 × 16 = 64 bytes. This allows a second group to load from a different PSSM row (but the same columns) within the same transaction, resulting in a two-way bank conflict. To resolve this, we employ two copies of the PSSM in shared memory. These copies are stored such that the i-th thread group consistently uses memory banks 0-15 if i is even, and banks 16-31 if i is odd. This ensures that accesses from different groups within the same transaction are directed to distinct memory banks, eliminating the bank conflict.

##### Per-cell computation and warp shuffles

Once the scores are loaded, threads perform computations for their assigned columns. Each thread performs the following operations for each cell: (1) adds the score from the scoring profile to the value from the previous diagonal cell, (2) sets the result to zero if it is negative, and (3) updates the local maximum score if the current cell’s value exceeds it. To facilitate data communication between threads within a group, particularly for diagonal dependencies, we use warp-shuffle operations. Since the group size is at most 32, all threads within a group belong to the same warp. Warp shuffles allow for efficient register-based data exchange between threads, avoiding slower shared memory access with explicit synchronization, or the even slower use of global memory.

##### Data permutation and vectorization

To double computational throughput, we utilize hardware instructions capable of processing two 16-bit values packed into a 32-bit word. However, direct packing of neighboring matrix columns into a single 32-bit word introduces dependencies that hinder independent processing. To address this challenge, we apply a data permutation technique that rearranges columns within a thread, aligning diagonal dependencies directly within the packed data. For example, in cases where a thread handles 32 columns, we pack columns (0, 16), (1, 17), (2, 18), and so forth. This arrangement ensures that each pair of packed columns corresponds to cells that are diagonally adjacent in the DP matrix. As a result, the dependency between packed values is restricted to the preceding value from the previous row, enabling full exploitation of vectorized operations for computational efficiency.

##### Long sequence handling

This strategy allows matrix tiles with up to 2,048 columns to be processed efficiently with a thread group of size 16. For protein sequences exceeding 2,048 residues, the profile matrix is divided into multiple tiles processed sequentially. In such cases, the last column of each tile is stored in global memory to serve as the starting point for the first column of the next tile.

##### Multi-GPU Parallelization

To maximize performance on systems with multiple GPUs, we distribute the reference database across the available GPUs. Specifically, the target database sequences are split into chunks, and each GPU processes a separate chunk against the query sequence. The results from each GPU are then combined to produce the final result list. This parallelization strategy reduces execution time and allows for processing larger databases by leveraging the combined available GPU memory.

##### Database streaming

We implemented several optimizations to increase processing speed and handle large datasets efficiently. We minimize host-GPU communication latency by partitioning the reference database into smaller batches, allowing processing to be pipelined via asynchronous CUDA streams. Here, data transfer of the next batch i + 1 from the host to GPU is overlapped with the processing of the current batch i on GPU. Notably, this approach allows us to efficiently handle databases larger than the available GPU memory.

Furthermore, to maximize GPU memory utilization and minimize data transfer, the workflow caches as many reference database batches as fit in GPU memory, and re-uses them for subsequent queries without re-transfer from the host. The streaming capability further extends to reading database batches directly from disk storage when the dataset exceeds available host RAM.

##### GPU server

MMseqs2 workflows are constructed through scripts that repeatedly invoke the `mmseqs` binary, each time specifying different modules to execute. We observed that each invocation requiring GPU resources incurs a CUDA initialization overhead of approximately 300ms. This startup cost can become substantial in complex workflows; for instance, during the ColabFold MSA search, the `ungappedprefilter` module is called six times (three iterative searches each against the UniRef30 and ColabFoldDB databases).

To circumvent this overhead, we introduced an optional, dedicated GPU server mode. Here, we launch a persistent background process that maintains the GPU context and becomes responsible for database caching and executes alignment computations. Subsequent `mmseqs ungappedprefilter` invocations communicate their requests to this running server process via Linux shared-memory, thereby avoiding the initialization penalty and reducing overall workflow execution time.

#### B.3. GPU-accelerated Smith-Waterman-Gotoh

In addition to the gapless filter on GPU, we implemented a version of Smith-Waterman-Gotoh with affine gap penalties operating on protein profiles (PSSMs) as a modification of CUDASW++4.0 (26) to align reference sequences to the same query profiles used in the filter. As described in the previous section, this required to transpose the computed DP matrix to place the profile along the x-axis, and leveraged the same parallelization strategies. Matrix tiles are processed by thread groups of size 4, 8, 16, or 32 using in-register computations with 32-bit capacity to avoid overflows.

In contrast to the filter algorithm, the Smith-Waterman-Gotoh algorithm does not allow for threads in the same group to operate on the same row in parallel since cells depend on their left neighbor. Consequently, a wavefront pattern is used to have threads work on different rows along the minor diagonals.

#### B.4. Cell Updates Per Second (CUPS)

Speed of dynamic programming (DP) algorithms is typically reported by converting runtime into the number of DP matrix cell updates that are performed per second; e.g., TCUPS (Trillions of Cell Updates Per Second) as TCUPS = (Σ_i_ m_i_ × n_i_)/(t × 10^12^), where t is the runtime in seconds and m_i_ and n_i_ are the lengths of the aligned sequences.

##### Efficiency analysis

We analyzed whether the proposed algorithm is able to effectively remove overheads incurred by memory accesses using half2 arithmetic by modeling the theoretical peak performance (TPP) of the utilized GPU hardware as:

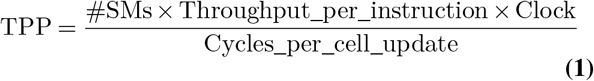

Where

- `Throughput_per_instruction` refers to the number of results per clock cycle per streaming multiprocessor (SM) of native arithmetic instructions on the considered hardware. A corresponding table for devices of various compute capabilities is provided in the CUDA documentation ^1^.
- `Cycles_per_cell_update` models the maximum attainable performance constrained by the algorithm structure and the specifications of the architecture. In our case, referring to the theoretical minimal number of clock cycles needed by an individual SM of the utilized GPU to calculate one DP matrix cell in M.

`Cycles_per_cell_update` can initially be determined by the inner loop in Algorithm 1 and thus set to three (i.e., two max instructions and one add instruction). However, as the L40S GPU used in most of our experiments enables dual ports add floating point operations, allowing to simultaneously issue add and `max` operations in a single SM cycle, `Cycles_per_cell_update` can be set to two (when using single-precision arithmetic) and to one (when using half2 arithmetic).

Disregarding the lookup operation to the PSSM, any value to register movements such as warp shuffles, any data transfers, and based on the specified `Throughput_per_instruction` of 64 for `max` instructions (the dependency bottleneck) on the L40S (compute capability 8.9), we can use Eq. 1 to calculate the `TPP` for half2 arithmetic on the L40S as follows:

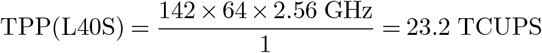

Provided the synthetic benchmark on a single L40S achieves a performance of up to 13.5 TCUPS (Fig. 1d), our approach is able to achieve an efficiency of 58% on an L40S GPU architecture, which shows that our optimizations, such as warp-shuffles and PSSM lookups from shared memory, are able to effectively transform the problem to compute-bound and minimize overheads from memory accesses.

### C. MMseqs2-GPU Workflow

The MMseqs2-GPU workflow starts with query and reference sets residing in CPU RAM. For the first step, i.e. filtering, query profiles are transferred to the GPU and permuted to enable efficient CUDA shared memory accesses during lookup. Additionally, the reference set is partitioned into smaller batches, which are transferred to the GPU. Throughout the filtering step, gapless alignment scores are stored for subsequent sorting. In the next stage, after all reference database batches have been processed, gapped, i.e. Smith-Waterman-Gotoh alignment scores are computed for top reference sequences satisfying inclusion thresholds using the same partitioned approach used for gapless alignment. Finally, the filtered reference sequences and their corresponding alignment scores are transferred back to the CPU.

### D. Hardware setup

With the exception of benchmarks executed on Google Colab compute, the same base system hardware setup was used. Specifically, the system featured two AMD EPYC 7742 64-core (2.25 GHz; Thermal Design Power (TDP) of 225W) CPUs (effectively 128 physical cores running 256 logical threads and TDP of 450W), with 1TB of DDR4 RAM and 2TB NVMe Intel SSDPE storage.

For the GPU benchmarks, the base system configuration additionally included either an L4, A100, L40S, or H100 PCIe NVIDIA GPU, which are set at TDP 72, 300, 350 and 350 Watts, respectively. The energy efficiency measurements additionally include a one socket EPYC 7313p 16-core (3 GHz; 155W TDP) system, with 256GB DDR4 RAM and L4 GPU.

### E. Benchmarks

#### E.1. Synthetic TCUPS benchmark

To measure TCUPS performance for MMseqs2-GPU (Fig. 1d), we performed a synthetic benchmark generating sequences of equal lengths for several lengths. For each possible DP matrix tile size l, a randomly generated query of length l was aligned to a database of five million randomly generated sequences of length l. Runtime was then converted to TCUPS. We executed the same benchmark using the CPU implementation of the gapless alignment described in “CPU-SIMD gapless filter”. We show speedup for each length l of the respective execution on 1, 2, 4 and 8 GPUs vs. the CPU execution. This synthetic benchmarks explores the best case scenario by employing uniform length, as this avoids reduced hardware utilization caused by adjacent CUDA thread groups processing different reference sequence lengths. For real-world database searches, the variable length effect is minimized by sorting the database sequences by length in ascending order.

#### E.2. Database scaling benchmark

To investigate performance characteristics of the MMseqs2-GPU `ungappedprefilter` implementation when the available GPU memory is exceeded, we measure TCUPs on real amino acid sequences based on the sequence sets described in section “Sensitivity”, with additional measurements conducted by extending the reference sequences 4, and 16 times (Fig. 1e). We extrapolated the runtime for the 16x replicated database on one L40S from a subset of 500 random queries out of all 6370 queries. TCUPS are similar for MMseqs2-GPU combined gapless and gapped executions for 1, 2, 4, 8, and 16 times the reference sequence set (Supplementary Figure 1).

#### E.3. Sensitivity

The same approach and datasets described in MMseqs2 (4) were used to conduct the sensitivity benchmark. This benchmark involved annotating full-length UniProt sequences with structural domain annotations from SCOP (37), designating 6,370 sequences as queries and 3.4 million as reference sequences. The full-length query sequences included disordered, low-complexity, and repeat regions, which are known to cause false-positive matches, especially in iterative profile searches. Additionally, 27 million reversed UniProt sequences were included as reference sequences (resulting in a total of 30,430,281 reference sequences).

Like in previous work, true-positive matches are defined as those with annotated SCOP domains from the same family, while false positives match reversed sequences or sequences with SCOP domains from different folds. The sensitivity of a single search is measured by the area under the curve (AUC) before the first false-positive match (ROC1), indicating the fraction of true-positive matches found with a better E-value than the first false-positive match. Sensitivity results for the various tools and modes, measured as ROC1, are summarized in the table below, with full benchmark details available in Supplementary Data 1.

| Tool | Mode | Sens. 1-it | 2-it | 3-it |
| --- | --- | --- | --- | --- |
| MMseqs2-CPU | -s 8.5 | 0.391 | 0.606 | 0.665 |
| MMseqs2-CPU | gapless | 0.4 | 0.612 | 0.669 |
| MMseqs2-GPU | gapless | 0.4 | 0.612 | 0.669 |
| BLAST |  | 0.401 |  |  |
| PSI-BLAST |  |  | 0.574 | 0.591 |
| JackHMMER |  | 0.4 | 0.614 | 0.685 |
| Diamond | ultra sens | 0.409 |  |  |

#### E.4. Speed

For speed benchmarks, the same query and reference sequences as for the sensitivity benchmarks were used, and methods’ parameters were set to reach comparable sensitivity where possible. Specifically, we ran the following methods:

- **JackHMMER v3.4** (2): at 1 iteration (equivalent to a phmmer search), with an E-value cutoff of 10, 000, using 128 threads (corresponding to the real number of cores, which differs from the actual number of threads), omitting the alignment section for the generated output.
- **BLAST v2.16.0** (1): with an E-value cutoff of 10, 000, using 128 threads, and limiting the number of filtered reference sequences to 4, 000.
- **PSI-BLAST v2.16.0** (1): at 2 iterations, with an E-value cutoff of 10, 000, using 128 threads, and limiting the number of filtered reference sequences to 4, 000.
- **Diamond ultra sensitive v2.1.9** (3): using `blastp`, the ultra-sensitive setting, with an E-value cutoff of 10, 000, using 128 threads, and limiting the number of filtered reference sequences to 4, 000.
- **MMseqs2-CPU k-mer commit 16-747c6** (4): using `easy-search` and the sensitivity at 8.5, with an E-value cutoff of 10, 000, using 128 threads, and limiting the number of filtered reference sequences to 4, 000.
- **MMseqs2-CPU gapless commit 16-747c6** (4): using `easy-search` and the prefilter mode at 1, with an E-value cutoff of 10, 000, using 128 threads, and limiting the number of filtered reference sequences to 4, 000.
- **MMseqs2-GPU commit 16-747c6:** using `easy-search`, the prefilter mode at 1 and enabling the GPU usage via the `--gpu 1 --gpu-server 1` options, with an E-value cutoff of 10, 000, using 1 thread, and limiting the number of filtered reference sequences to 4, 000.

We measured the total execution time of these methods based on the invocations above, and derived an average per-sequence execution. Additionally, we set up searches in either single batch mode, emulating the behavior in e.g., a protein 3D structure prediction server, or by batching sequences into groups of 10 and 100 sequences, or one batch of 6370 sequences, emulating the annotation of small protein sets or proteomes. Due to the prohibitive execution time of JackHMMER (2) in any batch mode, or Diamond (3) in single-batch mode, we reduced the query set to only 10%, or 637 query sequences to extrapolate results for the remaining cases. As batch 6370 would require to run 100% of the query set, and batching results from 1 to 100 for 10% of the query set showed negligible effect for JackHMMER, we excluded running this final benchmark with sustainability in mind.

#### E.5. Diamond scaling

Diamond (3) was developed to annotate vast metagenomic databases, and its performance is tuned to large query sets and batch sizes. In order to allow a fair comparison of Diamond’s performance, we thus performed a query database and batch scaling benchmark exclusively for Diamond. For this benchmark, all previous execution parameters were retained with the exception of the query database, since up-sampling the 6370 query set to obtain larger query database sizes is not a realistic setting. Instead, we randomly sampled (without replacement) from UniRef90 (38) to obtain query databases of size 100, 1,000, 10,000 and 100,000 sequences. We set the batch size equal to the database size for each run.

#### E.6. Structure prediction

To compare total runtime and the effect that MSA choice has on structure prediction accuracy, we replicated the ColabFold (28) benchmarks to predict the CASP14 (30) free-modeling queries (FM).

We measured MSA input generation and model inference times for the three configurations listed below. For all three configurations, we omit template search and relaxation and execute the default three recycling iterations.

- **AlphaFold2:** a baseline-configuration using the AlphaFold2 (11) pipeline, utilizing JackHMMER (2) and HHblits (21) to compute input MSAs from three databases, and structure prediction inference using AlphaFold2 weights. Specifically, this configuration utilizes the following homology search databases: UniRef90 2020_01 (JackHMMer, 139M sequences), MGnify 2018_12 (JackHMMer, 287M sequences), Uniclust30 2018_08 (HHblits, 15M profiles and 124M total), BFD first release (HHblits, 66M profiles and 2.1B total). Five AlphaFold models were run on one L40S GPU via the docker container from github.com/googledeepmind/alphafold (commit f251de6).
- **ColabFold MMseqs2-CPU:** a configuration using ColabFold v1.5.5, utilising MMseqs2 (commit 22115b) with default k-mer on CPU to compute input MSAs, and structure prediction inference loading AlphaFold2 weights. By default ColabFold utilizes recent versions of the databases, however, we retrieved older versions to approximate a fair comparison to the AlphaFold2 configuration, specifically UniRef30 2021_03 (29M representatives and 277M total) and ColabFoldDB 2021_08 (209M representatives and 739M total). Five AlphaFold models were run on one L40S GPU using ColabFold code obtained from github.com/sokrypton/ColabFold (commit 09993a8).
- **ColabFold MMseqs2-GPU:** a configuration using ColabFold v1.5.5, utilising MMseqs2-GPU (commit 22115b) with default gapless running on one L40S GPU to compute input MSAs, and structure prediction inference loading AlphaFold2 weights. We used the same databases and codebase as in the ColabFold MMseqs2-CPU configuration. All five models were run on one L40S GPU.

Structure prediction inference in all three configurations leveraged an L40S. To compare structure prediction accuracy, we computed several metrics between the predicted and ground truth structures, and focused results on template modeling score (TM-score; 39).

#### E.7. Cloud cost

We compared cloud cost to run the homology search methods presented in the speed benchmark. To do so, we retrieved on-demand hourly cost for AWS EC2 instances (retrieved on 2024/09/24) that best matched our hardware setup for instances in the US East (Ohio) region. We obtained:

- **CPU-based methods:** for JackHMMER, Diamond (ultra sensitive), MMseqs2-CPU k-mer, and BLAST, we selected the instance “c7a.32xlarge” with 128 physical CPU cores and 256 GBs of system RAM at an hourly cost of $6.57.
- **GPU-based method:** for MMseqs2-GPU, we selected the instance “g6e.xlarge” with 2 physical CPU cores and 32 GB of system RAM and one L40S GPU at an hourly cost of $1.86. For cloud costs utilizing 2 and 4 L40S, we selected the instance “g6e.12xlarge” with 24 physical CPU cores and 384 GB of system RAM and 4 L40S GPUs at an hourly cost of $10.493. For cloud cost calculations utilizing 8 L40S, we selected the instance “g6e.48xlarge” with 96 physical CPU cores and 1536 GB of system RAM and 8 L40S GPUs at an hourly cost of $30.131.

To obtain per-query-cost, we multiplied the hourly cost by the actual total runtime for each method and divided by the number of queries. All results can be found in Supplementary Material 1, “Cloud cost estimates”. Note that prices of cloud instances can differ regionally, and over time. To attempt a robust comparison, we reported the cost factor (i.e., the difference in cost compared to a baseline, which we selected to be MMseqs2-CPU k-mer) rather than the total theoretical cost.

#### E.8. Energy consumption

To measure average power utilization, we leveraged `powerstat` and `nvidia-smi` which allow to sample hardware counters (Running Average Power Limit; RAPL) included in CPUs and GPUs, and their memory. We performed this measurement for the full query set in single-batch mode for various methods. We then multiplied the power average by the execution times to obtain the total energy consumption.

#### E.9. Foldseek

We benchmarked Foldseek (31) using MMseqs2-GPU by sampling 6370 protein structures represented as 3Di strings (avg. length: 261 3Di letters). These were sampled from AFDB50 v4 (53.6M entries, avg. length: 264 3Di letters) against the same database. The database was indexed using createindex and stored in memory.

- **Foldseek-CPU k-mer commit c438b9** (31): using `search` with option `--db-load-mode 2` for fast index reading and 128 threads `--threads 128`.
- **Foldseek-GPU commit c438b9:** using `search` with option `--db-load-mode 2` for fast index reading, 128 threads `--threads 128` and `--gpu 1` for gapless GPU alignment.

We retrieved the Foldseek SCOPe-based sensitivity benchmark from github.com/steineggerlab/foldseek-analysis (commit 1737c71). Foldseek-GPU was executed with parameters `--max-seqs 2000 -e 10 --gpu 1`. Foldseek-CPU k-mer was executed with parameters `-s 9.5 --max-seqs 2000 -e 10`.

#### E.10. Colab benchmark

To compare the speed of MMseqs2-GPU to JackHMMER on more typically encountered hardware setup, we chose a paid Google Colab Pro environment with a 6-core/12-thread CPU, 64GB of system RAM and a NVIDIA L4 GPU with 24 GB RAM. We searched the same 20 CASP14 sequences as in section “Structure prediction” against the UniRef90 2022_01 (containing 144M proteins; benchmark performed on October 6th, 2024). As the reference database is 48.6 GB large, it does not fully fit into GPU memory (24GBs), leveraging system RAM streaming. We chose the following parameters for the search:

- **JackHMMER v3.4:** at 1 iteration (equivalent to a ph-mmer search), with an E-value cutoff of 10, 000, using 12 threads and omitting the alignment section for the generated output.
- **MMseqs2-GPU commit 81ddab:** using `easy-search`, enabling the GPU usage via the `–gpu 1` options, with an E-value cutoff of 10, 000, using 12 thread, and limiting the number of filtered reference sequences to 4, 000.

## Acknowledgments

We thank Yuxing Peng, Maximilian Stadler, and Maximilian Baust for invaluable suggestions and support, Arne Elofsson and Shiraz A. Shah for reporting issues and providing feedback, Eli Levy Karin, and the anonymous RECOMB-seq reviewers for critical feedback on the manuscript. M.S. acknowledges support by the National Research Foundation of Korea grants (2020M3-A9G7-103933, 2021-R1C1-C102065 and 2021-M3A9-I4021220, RS-2024-00396026), Samsung DS research fund, Creative-Pioneering Researchers Program and Novo Nordisk Foundation (NNF24SA0092560). M.M. acknowledges support from the National Research Foundation of Korea (grant RS-2023-00250470). B.S. acknowledges support from the Deutsche Forschungsgemeinschaft (DFG, German Research Foundation) – project number 439669440 TRR319 RMaP TP C01. The funding body did not participate in the design of the study and collection, analysis, and interpretation of data and in writing the manuscript.

## Author contributions

**Felix Kallenborn:** Software; Writing - Original Draft; Writing - Review & Editing. **Alejandro Chacon:** Software; Investigation, Writing - Original Draft; Writing - Review & Editing. **Christian Hundt:** Resources; Project administration. **Hassan Sirelkhatim:** Visualization; Investigation; Validation. **Kieran Didi:** Software; Investigation. **Sooyoung Cha:** Software; Investigation. **Christian Dallago:** Conceptualization; Resources; Writing - Original Draft; Writing - Review & Editing; Supervision; Visualization; Project administration. **Milot Mirdita:** Conceptualization; Methodology; Software, Investigation; Writing - Original Draft; Writing - Review & Editing; Supervision. **Bertil Schmidt:** Software; Methodology; Conceptualization; Resources; Writing - Original Draft; Writing - Review & Editing; Supervision;. **Martin Steinegger:** Software; Methodology; Conceptualization; Resources; Writing - Original Draft; Writing - Review & Editing; Supervision.

## Data availability

Data utilized to perform benchmarks for this study are freely available. For speed, sensitivity, and energy consumption benchmarks, we leveraged target sequences stored at mmseqs.steineggerlab.workers.dev/targetdb.fasta.gz and reference sequences stored at mm- seqs.steineggerlab.workers.dev/query.fasta. For the folding benchmark, CASP14 targets are available at pre- dictioncenter.org/casp14/index.cgi, while the reference ColabFold databases are available at colabfold.mmseqs.com. For Foldseek benchmarks, we retrieved data and scripts from github.com/steineggerlab/foldseek-analysis.

## Code availability

All code developed in this study is available under MIT license and documented at mmseqs.com. Analysis scripts are available at github.com/steineggerlab/mmseqs2-gpu-analysis.

## Supplementary materials

**Supplementary Figure 1.**
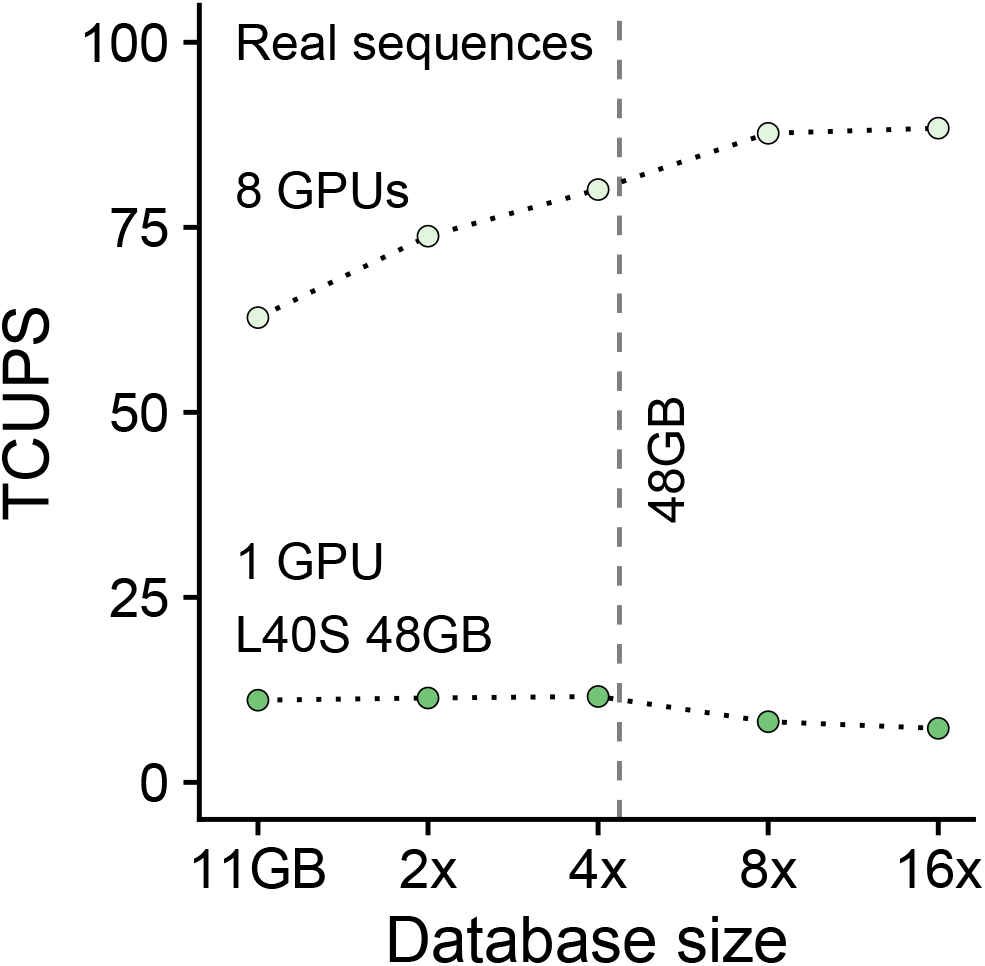
Combined gapless and gapped alignment TCUPS. TCUPS of 1 and 8 GPU executions of the combined MMseqs2-GPU gapless and gapped alignment workflow for 6370 queries against target sets of 1, 2, 4, 8, and 16 times a 30M protein database (Methods “Sensitivity”). 8 and 16 times executions exceeds GPU RAM and are processed with database streaming. The latter is processed with 7.3 TCUPS/11.6 TCUPS ≈ 63 % of in-memory processing speed.

## Footnotes

1 https://docs.nvidia.com/cuda/cuda-c-programming-guide/index.html#arithmetic-instructions, Accessed May 12th, 2025.

