## Supplementary Figure 1 for "GPU-accelerated homology search with MMseqs2"

### Supplementary materials

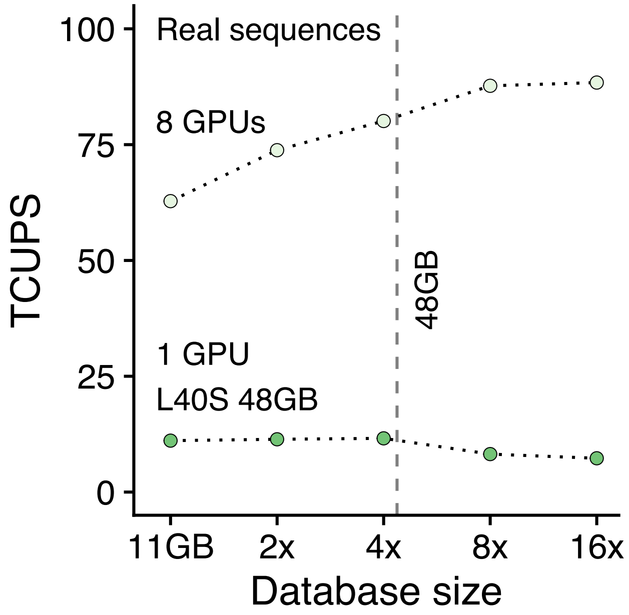


**Supplementary Figure 1. Combined gapless and gapped alignment TCUPS.** TCUPS of 1 and 8 GPU executions of the combined MMseqs2-GPU gapless and gapped alignment workflow for 6370 queries against target sets of 1, 2, 4, 8, and 16 times a 30M protein database (Methods "Sensitivity"). 8 and 16 times executions exceeds GPU RAM and are processed with database streaming. The latter is processed with $7.3\,\mathrm{TCUPS}/11.6\,\mathrm{TCUPS} \approx 63\%$ of in-memory processing speed.
